# Cerebro-spinal somatotopic organization uncovered through functional connectivity mapping

**DOI:** 10.1101/2024.04.11.588866

**Authors:** Caroline Landelle, Nawal Kinany, Benjamin De Leener, Nicholas D. Murphy, Ovidiu Lungu, Véronique Marchand-Pauvert, Dimitri Van De Ville, Julien Doyon

**Author notes:** These authors share first authorship (Caroline Landelle, Nawal Kinany). These authors share senior authorship (Dimitri Van De Ville, Julien Doyon).

## Abstract

Somatotopy, the topographical arrangement of sensorimotor pathways corresponding to distinct body parts, is a fundamental feature of the human central nervous system (CNS). Traditionally, investigations into brain and spinal cord somatotopy have been conducted independently, primarily utilizing body stimulations or movements. To date, however, no study has probed the somatotopic arrangement of cerebro-spinal functional connections *in vivo* in humans. In this study, we used simultaneous brain and cervical spinal cord functional magnetic resonance imaging (fMRI) to demonstrate how the coordinated activities of these two CNS levels at rest can reveal their shared somatotopy. Using functional connectivity analyses, we mapped preferential correlation patterns between each spinal cord segment and distinct brain regions, revealing a somatotopic gradient within the cortical sensorimotor network. We then validated this large-scale somatotopic organization through a complementary data-driven analysis, where we effectively identified spinal cord segments through the connectivity profiles of their voxels with the sensorimotor cortex. These findings underscore the potential of resting-state cerebro-spinal cord fMRI to probe the large-scale organization of the human sensorimotor system with minimal experimental burden, holding promise for gaining a more comprehensive understanding of normal and impaired somatosensory-motor functions.

## 1. Introduction

Topographic organization, characterized by an orderly mapping of neural representations for distinct sensory inputs or motor outputs, is a fundamental and ubiquitous feature of the central nervous system (CNS) (Patel et al. 2014; Eickhoff et al. 2018). A quintessential example is the somatotopic arrangement observed in the somatosensory and motor cortices, where distinct neural patterns are linked to perception and movement of specific body parts. This arrangement is classically illustrated using Penfield’s homunculi (Penfield & Boldrey, 1937), depicting lower-limbs near the midline, the face laterally in the cortex, and a large representation of the hands in between. Although the reality may be more nuanced, with overlapping and intermingled representations among body parts (Gordon et al., 2023; Meier et al., 2008; Muret et al., 2022), the relationship between cerebral organization and body representation remains widely accepted. Besides the cortex, this somatotopic organization is also evident throughout the sensorimotor hierarchy. At the core of the brain-body axis, the spinal cord mirrors the body’s representation with pools of neurons arranged into rostro-caudal segments, innervating distinct dermatomes and myotomes from head to toe (ten Donkelaar, 2011a, 2011b)

Mapping somatotopy has been a longstanding endeavor in neuroscience, both to understand the healthy human sensorimotor system and to explore plastic changes associated with age or injuries (Muret & Makin, 2021; Newbold et al., 2020). One avenue has entailed the intraoperative application of electrical current directly to the cortex (Desmurget & Sirigu, 2015; Penfield & Boldrey, 1937; Roux et al., 2018, 2020) or spinal nerves (Keegan & Garrett, 1948; Schirmer et al., 2011). However, the invasive nature of these methods significantly limits the scope of their application. Conversely, functional magnetic resonance imaging (fMRI) has emerged as a powerful non-invasive alternative, extensively utilized to delineate limb representation through motor tasks and somatosensory stimulations, in both cortical and subcortical brain areas (Boillat et al., 2020; Gordon et al., 2023; Karbasforoushan et al., 2022; Tal et al., 2017; Zeharia et al., 2015). In the spinal cord, these investigations have been more limited, even though recent progress in spinal cord fMRI (see Landelle et al. 2021; Kinany et al. 2023a for reviews) have paved the way for a handful of studies mapping activity patterns related to perception or movement of distinct body parts (Kinany et al., 2019; Lawrence et al., 2008; Weber et al., 2020). Besides, recent research has shown that the segmental arrangement of the spinal cord could be mapped at rest using spontaneous activity and data-driven network extraction methods (Kinany et al., 2024). Overall, task-based paradigms have been the prevailing approach to study body mapping along the neural axis. While these protocols have been essential for advancing our understanding of somatotopy, they face obstacles such as difficulties in accessing or moving specific body parts. In addition, they can be challenging to deploy in clinical environments due to factors like participant inability to perform tasks, equipment constraints, and experiment duration.

Despite evidence indicating shared functional organizational principles between the brain and spinal cord (ten Donkelaar, 2011a, 2011b; Usuda et al., 2022), these regions have traditionally been studied as distinct entities. This segregated approach has posed limitations on mapping the somatotopic organization of sensory and motor pathways along the CNS axis. Accordingly, adopting a more holistic approach is particularly relevant for gaining a more comprehensive understanding of normal and impaired somatosensory-motor functions. Only very recently, advances in fMRI have enabled the simultaneous investigation of both brain and spinal cord functional activity across the sensorimotor hierarchy (see Kaptan et al. 2024; Tinnermann et al. 2021a for reviews). Despite the potential of cerebro-spinal fMRI, the literature on this topic remains sparse, with most studies focusing on pain (Sprenger et al. 2015, 2018; Oliva et al. 2022; Tinnermann et al. 2017, 2021b, 2022) and a limited number on motor tasks (Vahdat et al. 2015; Khatibi et al. 2022; Braaß et al.2023; Kinany et al. 2023b). To the best of our knowledge, only one study has deployed simultaneous cerebro-spinal fMRI to investigate the large-scale functional organization of brain and spinal cord activity, using resting-state recordings (Vahdat et al., 2020). By averaging signals across multiple segments of the cervical spinal cord and examining their correlations with the brain, this work offered first insights into the unified functional organization of spontaneous activity within the CNS. Specifically, the observed patterns of correlations aligned with established anatomical principles related to lateralization and sensory-motor subdivisions, underscoring the functional relevance of cerebro-spinal intrinsic activity. Based on such findings, it is thus possible to posit that the spontaneous activity between the spinal cord and the sensorimotor cortex also follows a somatotopic organization principle, reflecting a topographical body representation along the CNS axis. To date, however, no study has probed the somatotopic arrangement of cerebro-spinal functional connections *in vivo* in humans.

To examine this hypothesis, we leveraged simultaneous brain-spinal cord resting-state fMRI combined with advanced functional analysis methods. Departing from the prevailing task-based approach in studying the topographic organization of the CNS, we proposed to harness the intricate interplay of spontaneous fluctuations and co-variations between the brain and spinal cord to uncover their shared somatotopic organization. First, we expected that cerebro-spinal functional connectivity analyses, using spinal cord functional levels as seeds, could unveil the brain’s somatotopic organization. Second, we sought to validate this shared somatotopic organization by assessing whether spinal cord functional levels can be identified, in a data-driven manner, based on their functional connectivity with sensorimotor cortical regions. With its versatility and low experimental burden, this approach to mapping somatotopic organization holds potential for both fundamental research and clinical applications.

## 2. Methods

### 2.1 Participants and MRI data acquisitions

Thirty-one right-handed healthy participants (16 females; age 32.8 ± 6.8 years old) were included in this study. Participants reported no neurological or sensorimotor disorders and had no contraindication for MRI. The experiment was approved by the local ethics committee (MUCH REB 2019-4626) and all participants gave their written consent in accordance with the Helsinki Declaration. MRI data were acquired using a 3-T MRI scanner (Magnetom-Prisma, Siemens, Erlangen, Germany) at The Neuro (Montreal Neurological Institute, Canada) using a 64-channel phase-array head coil (1-7 elements active) and a neck coil (1-2 elements active). Participants were placed in the scanner in supine position, while wearing Neck and Brachial Plexus SatPads. Throughout the scanning session, the participants were instructed to relax, to minimize motion, and to swallow gently when needed.

Anatomical images were acquired using a high-resolution T1-weighted sequence that covered the whole brain and the cervical spinal cord down to the T1 vertebrae (Repetition time (TR)/Echo Time (TE) = 2300/3.3 ms, MPRAGE sequence, generalized autocalibrating partially parallel acquisition (GRAPPA) with integrated parallel acquisition technique (iPAT) = 2, flip angle = 9°, resolution = 1.3 x 1.3 x 1.3 mm^3^, transversal acquisition, Field of view (FOV) = 228 × 364 × 375 mm^3^ LPI orientation).

To achieve continuous coverage of the brain and spinal cord, we devised a novel fMRI acquisition protocol, enabling simultaneous imaging of both regions within a single field-of-view (FOV). Specifically, functional images were acquired using a multiband gradient-echo EPI sequence covering the brain and cervical spinal cord (TR/TE = 1550/23 ms, axial FOV = 192 x 192 mm^2^, generalized autocalibrating partially parallel acquisition (GRAPPA) with integrated parallel acquisition technique (iPAT) acceleration factor for phase encoding direction = 2 and multiband factor for slice encoding direction acceleration factor = 3, flip angle = 70°, in-plane resolution = 1.6 x 1.6 mm^2^, slice thickness = 4 mm, transversal acquisition, number of slices = 69, number of volumes = 230, duration = 6 min). The slices were positioned using the high-resolution T1w image so that the FOV was placed parallel to the spinal cord. The shim volume was manually set to focus on the spinal cord. Examples of raw T1w and functional images are available in Fig. S1.

The functional data were acquired during resting state (*i*.*e*., no explicit task) with participants instructed to refrain from specific thoughts and focus on observing the *Inscapes* video (Vanderwal et al., 2015). Physiological recordings were acquired using a pulse sensor and a respiration belt (Siemens Physiology Monitoring Unit).

### 2.2 Processing

All acquired images were sorted (with dcm2niix v1.0.20181125), transformed in NIFTI and stored using the Brain Imaging Data Structure (BIDS) standard (with dcm2bids v2.1.4). The functional and structural images were processed using in-house python pipelines based on the Spinal Cord Toolbox (SCT, version 5.6.0) (De Leener et al., 2017) the Oxford Center for fMRI of the Software Library (FSL, version 5.0), the Statistical Parametric Mapping (SPM12, running on Matlab 2021b), the Tapas PhysiO toolbox (release 2022a, V8.1.0) (Kasper et al., 2017), and the Nilearn toolbox (version 0.9.1).

#### 2.2.1 Preprocessing

First, slice-timing correction was applied to the functional images, followed by cropping of the functional and anatomical images to perform tailored preprocessing on the brain and spinal cord. The preprocessing included the following steps:

i) *Motion correction of functional images*. Spinal cord motion correction was applied using slice-wise realignment and spline interpolation (with SCT, *sct_fmri_moco*) inside a 30mm cylindrical mask centered on the spinal cord’s centerline and covering the gray matter (GM), white matter (WM), and cerebrospinal fluid (CSF). Brain motion correction was performed using rigid-body realignment (with FSL, MCFLIRT, Jenkinson et al., 2002) after removing the non-brain tissues (with FSL, BET, Smith, 2002). We quantified motion between two consecutive volumes by calculating framewise displacement (FD) using realignment motion parameters at both brain and spinal cord levels. The mean spinal cord FD (averaged over the x and y axis) was 0.11 ± 0.04 mm and the mean FD for the brain was 0.063 ± 0.02 mm (Fig. S2a). None of our participants exceeded the predetermined threshold for excessive motion (*i*.*e*., FD_brain_ or FD_spinalcord_ > 0.3 mm, Landelle et al., 2023). Following motion correction, the temporal signal-to-noise ratio (tSNR) was computed to evaluate the quality of the functional signals over time. The tSNR was obtained for each voxel by dividing the mean signal intensity across time by its standard deviation. At the group level, we obtained a mean tSNR of 21.68 ± 2.63 within the spinal cord and 46.20 ± 3.8 within the brain (Fig. S2b-c).
ii) *Segmentation of functional and structural images*. Spinal cord segmentation (GM+WM) was first performed automatically on the T1w spinal cord anatomical images (with SCT, *sct_propseg*). Segmentation of the mean motion corrected spinal functional images was done using a two-step process: the spinal centerline was extracted manually and subsequently the cord (GM+WM) and the CSF were extracted (with SCT, *sct_propseg*). Spinal cord tissue segmentations were visually inspected and manual adjustments were performed when necessary. Brain tissues segmentation was performed automatically on T1w brain anatomical images (with CAT12, an SMP12 extension).
iii) *Time series denoising of functional images* (see details in Time series denoising).
iv) *Normalization into PAM50 or MNI template*. The normalization of the spinal cord functional images was done in two steps. First, the T1w image was warped into the PAM50 space (0.5 x 0.5 x 0.5 mm^3^) using cord segmentation and disc labeling (with SCT, *sct_register_to_template*). Second, the mean functional image was warped into the PAM50 space using the warping field obtained at the previous step (*i*.*e*., from T1w to PAM50 space) to initialize the registration (with SCT, *sct_register_multimodal*). The brain normalization was done in three steps using SPM12. First, the mean functional image was coregistered to the T1w space. Second, the T1w image was warped into the MNI template (2 x 2 x 2 mm^3^). Third, the resulting warping field (*i*.*e*., from T1w to MNI space) was applied to the brain functional images in T1w space.
vi) *Spatial smoothing*. The denoised and normalized images were subsequently smoothed using a 3D Gaussian kernel with a full width half maximum (FWHM) of 3 x 3 x 6 mm^3^ for the spinal cord and 6 x 6 x 6 mm^3^ for the brain (with nilearn, *nilearn*.*image*.*smooth_img*).

#### 2.2.2 Time series denoising

To ensure consistency and minimize the risk of introducing artifactual variations in the brain and spinal cord signals, identical sets of regressors were used to denoise the motion-corrected time series of both regions. For each participant, we modeled nuisance regressors to account for physiological noise (Tapas PhysiO toolbox, an SPM extension, Kasper et al., 2017). First, we computed noise regressors from peripheral physiological recordings (respiration and heart rate) using the RETROspective Image CORrection (RETROICOR) procedure (Glover et al., 2000). Specifically, we modeled three cardiac and four respiratory harmonics, and one multiplicative term for the interactions between respiratory and cardiac noise (18 regressors in total, similar to Kinany et al., 2024; Tinnermann et al., 2017). Second, we used the CompCor approach (Behzadi et al., 2007) to identify non-neural fluctuations by extracting the first principal components of the unsmoothed brain or spinal cord cerebrospinal fluid (CSF) signal in the participant’s native space (12 components for the brain and 5 components for the spinal cord). The first five discrete cosine transform (DCT) basis functions were added for detrending. These nuisance regressors were finally combined with the six brain and two spinal cord motion parameters. The removal of the noise confounds was based on a projection on the orthogonal of the fMRI time-series space and was applied orthogonally to a band-pass temporal filter (0.01-0.17Hz, with Nilearn, *img*.*clean_img*). Note that no temporal filter was applied for iCAP analyses (see Section 2.2.3), as the hemodynamic deconvolution enables the use of the full-spectrum fMRI signal (Karahanoğlu & Van De Ville, 2015). After applying (smoothed data) or not (unsmoothed data) the spatial smoothing, the denoised signal was demeaned and standardized.

#### 2.2.3 Functional connectivity analyses

##### Innovation-driven co-activation patterns (iCAPs)

We employed the iCAP framework to uncover brain (Karahanoğlu & Van De Ville, 2015) and spinal cord (Kinany et al., 2024) networks. This approach allows us to identify relevant functional regions of interest specific to our population, in a data-driven manner. Briefly, the iCAP framework incorporates two main features: *i)* the extraction of transient activity using Total Activation (TA, (Karahanoğlu et al., 2013)) and *ii)* the temporal clustering of these signals. Specifically, TA was first used to extract activity-inducing signals from the smoothed denoised time series, using a regularized hemodynamic deconvolution. Transient activity (also called innovation signals) was obtained as the temporal derivative of these activity-inducing time courses. Finally, frames with similar and significant transitioning activities were K-means clustered to obtain group-level iCAP maps. For the spinal cord, the value of K was set to match the number of segmental levels within the field-of-view, *i*.*e*., K = 7 (Frostell et al., 2016; Kinany et al., 2024). For the brain, we opted for K = 10, to delineate the core resting-state networks. In both instances, consensus clustering was employed to validate the stability of the networks obtained for the selected K values (Monti et al., 2003)

##### Seed-to-voxels analysis

Seeds were defined using the gray matter of the seven spinal cord iCAPs. At the individual level, the unsmoothed time series within each of the seven seeds were extracted and averaged. The functional connectivity (FC) strength between the average spinal time series and gray matter brain voxels was assessed using Pearson’s correlation. This process generated seven brain correlation maps for each individual, which were converted using Fisher’s r-to-z transformation. These participant-specific maps were then averaged across individuals to yield seven group-level FC maps. Subsequently, a “winner-take-all” selection method was used on these FC maps to assign, for each brain voxel within the iCAP-derived sensorimotor cortex (SMC) network, the spinal functional level to which it exhibited the strongest FC. This enabled the assignment of each brain voxel’s preferential FC to one of the seven spinal functional levels, beyond all other levels. This technique is powerful to reveal gradients of selectivity for topographic maps and is commonly employed to uncover somatotopic representation (Gordon et al., 2023).

In order to assess the reproducibility across participants, winner-take-all maps were also computed at the individual level. Then, within the seven group-level winner-take-all masks (each corresponding to one spinal segmental level), the number of voxels assigned to each spinal level was calculated for each individual. Statistical comparisons were computed for each of the seven group-level masks by fitting the resulting number of voxels in a linear mixed model using the R package ‘lme4’. This model took into account the effects of spinal level assignment (fixed-effect), the variability between participants (random effects) and the residual error.

##### Task-related analysis

An additional publicly accessible fMRI dataset was used to capture brain somatotopic organization related to body movements (Ma et al., 2022). This brain fMRI dataset was specifically designed to explore whole-body somatotopic mapping in a cohort of 61 healthy adults (33 females; age 22.8 ± 2.3 years old), focusing on bilateral movements of various body parts: including toes, ankles, legs, upper arms, forearms, wrists, fingers, jaws, lips, tongue, and eyes. Two runs were performed for each movement. Importantly, we only included jaw, upper arm, forearm, wrist and finger movements for further analyses, as face movements and lower limb movements are respectively innervated by cranial and lumbosacral nerves (Keegan & Garrett, 1948; Schirmer et al., 2011). The authors provide denoised beta maps for each participant, combining the two runs using a fixed-effects model. We conducted group-level analyses for each movement of interest using a one-sample t-test with beta images from all participants. The resulting p-maps were Z-scored and thresholded at Z = 5 (uncorrected p<0.000001). Additionally, a winner-take-all map illustrating preferential sensorimotor activity for these five conditions was computed.

##### Connectivity-based parcellation

In order to parcellate the cervical spinal cord in a data-driven manner, using its interaction with the brain as input, we conducted a FC-based parcellation analysis (Eickhoff et al., 2015). Specifically, our objective was to investigate the feasibility of segmenting the spinal cord into its functional levels, based on its connectivity with sensorimotor regions in the brain.

i. *Extraction of similarity matrices* - For each participant, Pearson correlation coefficients were computed between every voxel within the spinal cord gray matter (extending from C1 to C7 segmental levels) and each voxel within the SMC network, as defined by the corresponding iCAP. These correlation coefficients were then converted using Fisher’s r-to-z transformation, yielding a correlation matrix (Z) of size Nvox_sc x Nvox_smc. To characterize the similarity between cerebro-spinal FC-profiles, we computed the functional similarity matrix as the cross-correlation of Z, resulting in a matrix of size Nvox_sc x Nvox_sc (Balsters et al., 2016; Johansen-Berg et al., 2004; Kim et al., 2010). These similarity matrices were averaged across participants for subsequent analysis.
ii. *Clustering of similarity matrices -* A hierarchical clustering algorithm (average linkage) (Johnson, 1967) was applied to cluster spinal voxels based on the dissimilarity of their FC profiles with the sensorimotor iCAP (*i*.*e*., 1 - average similarity matrix). The resulting dendrogram can be cut at different levels to yield clusters of varying granularity. Aligned with our objective of recovering the spinal functional levels corresponding to the seven segmental levels included in our study (Frostell et al., 2016), we cut the dendrogram at K = 7, but also examined neighboring solutions (±2 around the target K). Dice coefficients were computed to assess the spatial agreement between the cluster maps of the 7-cluster solution and a reference atlas of segmental levels (Frostell et al., 2016), matched using Maximum weight matching (Crouse, 2016).
iii. *Stability across participants -* To assess the stability of the parcellation procedure across participants, two analyses were done. First, we evaluated the correlation between the similarity profiles of each pair of participants for every spinal cord voxel. These voxelwise maps were averaged to obtain a single map representing the mean stability across all individuals. Second, we applied the hierarchical clustering procedure separately to individual similarity matrices, also segmenting the dendrogram into seven clusters. To evaluate the agreement between individual and group labels, we computed a contingency matrix between the two sets of labels for each participant. Values were normalized by the number of voxels in each cluster to mitigate bias towards larger clusters. This matrix was utilized to align individual labels with those from the group-level clustering, ensuring each individual label corresponded to at most one group-level label. Heatmaps of the individual clusters were generated by summing the binary maps corresponding to each label across all participants.
iv. *Connectivity of spinal FC-based segments -* To visualize the FC profiles of each spinal cluster, we computed the mean brain FC map corresponding to all voxels within each cluster index for each participant (*i*.*e*., seven maps per participant). These participant-specific maps were then averaged across individuals, in order to obtain the mean brain connectivity profile for each cluster index. A winner-take-all analysis was then applied to generate a map highlighting preferential FC patterns between the clusters and brain voxels.

## 3. Results

### 3.1 Identifying brain and spinal cord sensorimotor networks

To identify sensorimotor networks in both the brain and spinal cord, we employed the iCAP data-driven framework (Karahanoğlu & Van De Ville, 2015). We uncovered in total 10 components for the brain, which enabled the extraction of a cortical SMC network, alongside other established brain resting-state networks (Figs. 1a and S3). Specifically, brain iCAP3 featured a sensorimotor cortical network including the bilateral precentral and postcentral gyrus, as well as the bilateral parietal operculum. The remaining 9 iCAPs delineated cerebellar, auditory, lateral visual, ventral default mode, median visual, right fronto-parietal, left fronto-parietal, dorsal default mode and dorsal attentional networks (Fig. S3). Consensus clustering underscored the high robustness of these networks, as evidenced by the consensus values observed across subsets of the data (average consensus = 0.84). In the spinal cord, we extracted seven components based on anatomical knowledge regarding the number of segmental levels covered in the dataset (Frostell et al., 2016). Similar to the brain, these networks exhibited remarkable stability, with an average consensus value of 0.93. The seven iCAPs demonstrated a clear rostro-caudal organization, largely aligning with segmental levels (Dice coefficients: C1 = 0.86; C2 = 0.68, C3 = 0.71; C4 =0.86; C5 = 0.86; C6 = 0.89; C7 = 0.93), and specific to our population. All components encompassed both dorsal (*i*.*e*., sensory) and ventral (*i*.*e*., motor) bilateral regions.

**Figure 1.**
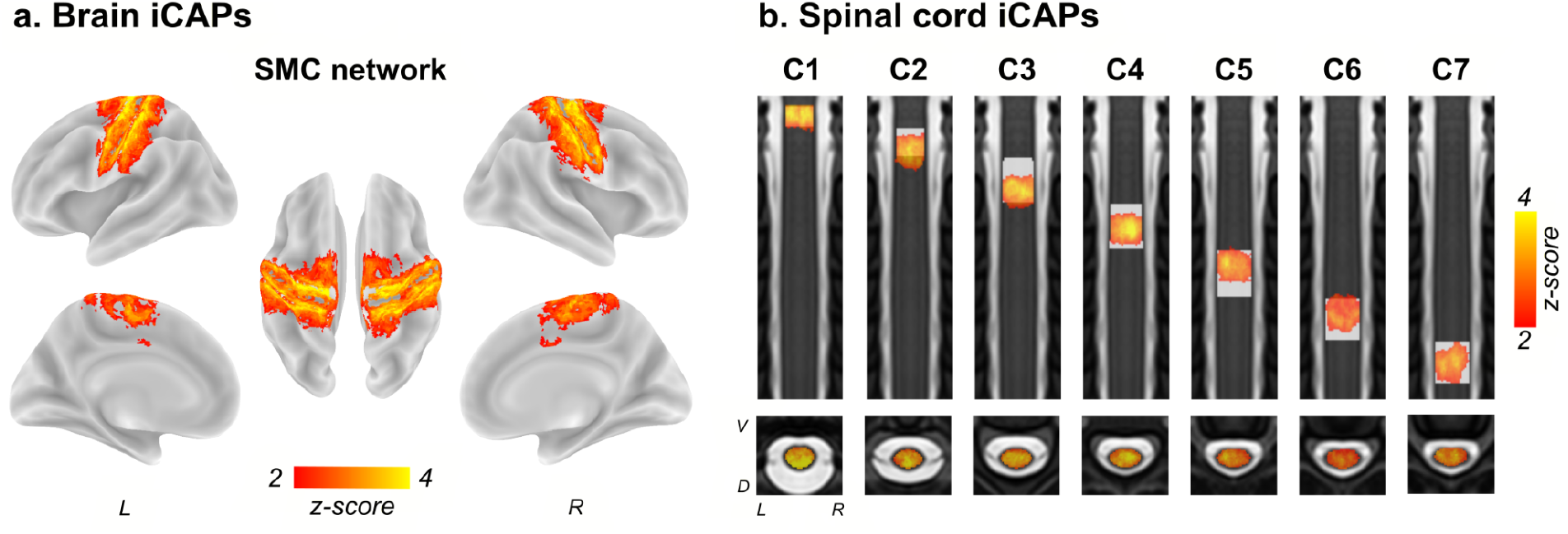
Spatial patterns of the sensorimotor networks at the brain and spinal cord levels. **a**. Brain innovation-driven co-activation patterns (iCAPs) corresponding to the sensorimotor cortex (SMC) network. The 9 other brain iCAPs are available in the supplementary material (Fig. S3). **b**. Spinal cord iCAPs presented in rostro-caudal order. The label of the iCAPs (from C1 to C7) are determined based on the spinal segmental atlas (Frostell et al., 2016), plotted as a white underlay. L: left; R: right; V: ventral; D: dorsal.

### 3.2 Somatotopic organization of spinal-level dependent functional connectivity within the SMC network

To assess the topographical organization of functional connectivity between the spinal cord and the brain we first conducted a seed-to-voxels correlation analysis. The seeds were defined as the gray matter of the seven spinal cord levels (*i*.*e*., the spinal functional levels obtained through the spinal iCAP analysis, Fig. 1b), while the target voxels were located within the SMC network (*i*.*e*., identified with the brain iCAP analysis, Fig. 1a). Then, a winner-take-all selection method was used to assign the winning seed, based on higher FC, to each voxel within the SMC network. This analysis revealed the topographical organization of cerebro-spinal FC across the SMC cortical surface, demonstrating a bilateral spinal level-dependent gradient (Fig. 2b). To better describe its spatial features, FC selectivity for distinct spinal levels is presented for the left hemisphere in Fig. 2c (see right hemisphere in Fig. S4). C1C2 levels connectivity exhibited an extended cluster in the premotor cortex (PMc). The order of progression from medial to lateral for other spinal levels along the SMC was approximately: C4, C5, C6, C7, a second representation for C4 and C5, and laterally the C3 level. In this gradient, the map corresponding to C7 had the largest spatial extent (number of voxels: C3 = 1808; C4 = 1079, C5 = 596, C6 = 1787, C7 = 2538). The preferential organization of the cerebro-spinal FC in the SMC was also evident when assessing the reproducibility in individual participants (Fig. S4, Table S1).

**Figure 2.**
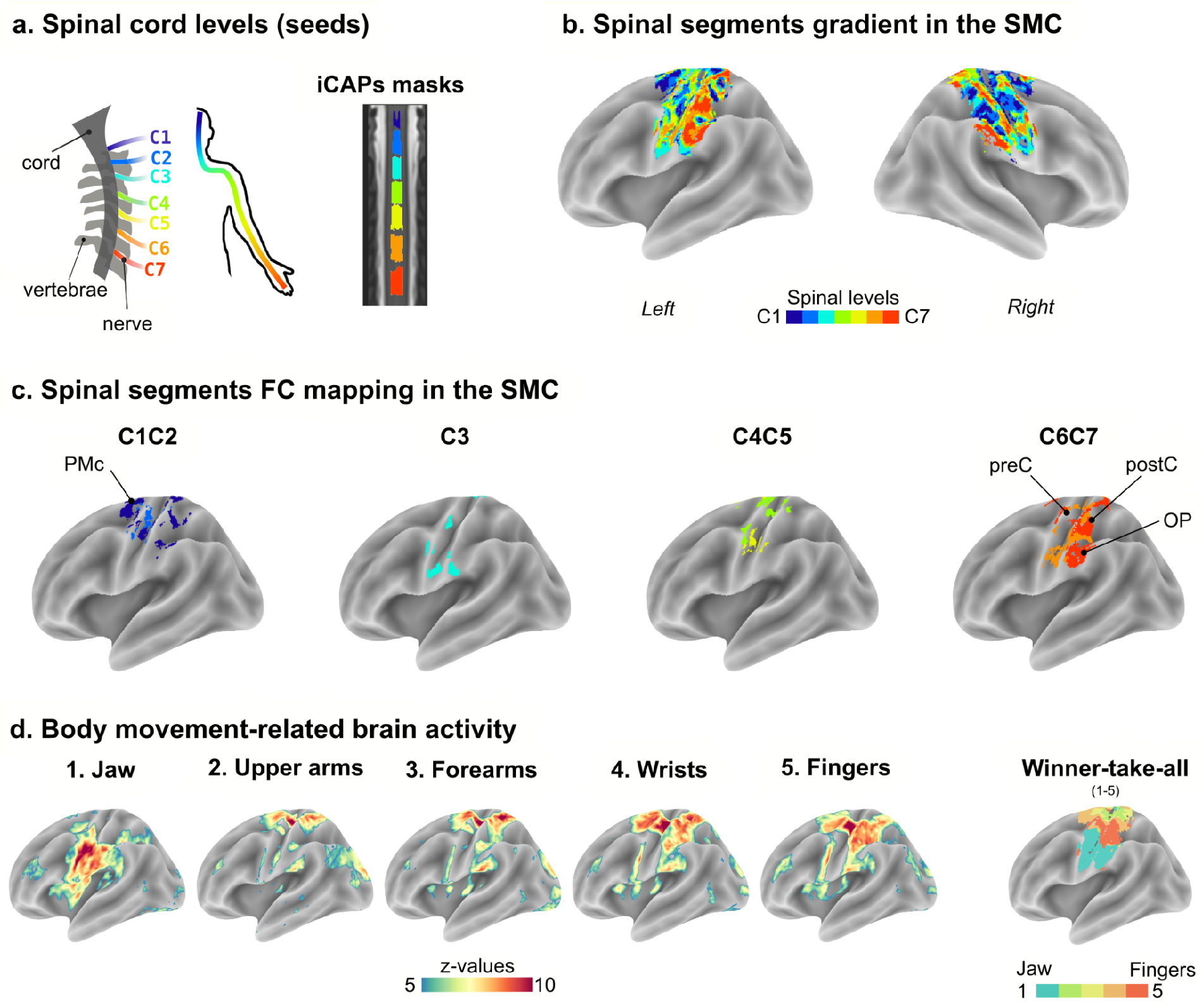
Gradient of cervical spinal cord functional connectivity along the SMC network. **a**. Spinal cord levels (seeds). Left: Schematic representation of the first seven cervical nerves, corresponding to the spinal segmental levels C1 to C7. Right: gray matter masks corresponding to the seven spinal cord iCAPs displayed on a coronal view of the PAM50-T2w template. **b**. Group level winner-take-all maps, smoothed (3mm^3^) for visualization purposes. Each color of the gradient indicates the FC selectivity for each iCAP-defined segment (C1: dark blue, C2: blue, C3: cyan, C4: green, C5: yellow, C6: orange, C7: red). **c**. The unsmoothed discrete winner-take-all map of the dominant hemisphere (left) was thresholded to display only one or two segments, simplifying the interpretation of spinal-level dependent FC within the SMC network. **d**. Group-level activation maps (Z-scores) for 61 participants, resulting from five bilateral movements of the upper body (jaw, upper arms, forearms, wrists and fingers movements). Statistical maps were thresholded at Z = 5 (p<0.000001 uncorrected). The last panel shows winner-take-all maps for these five conditions, in our sensorimotor iCAP mask. Task-related data were derived from a publicly available dataset (Ma et al., 2022). SMC: sensorimotor cortex; PMc: premotor cortex; preC: pre-central gyrus; postC: post-central gyrus; OP: parietal operculum

An additional dataset (Ma et al., 2022), including data for five bilateral movement conditions involving the upper body (jaw, forearms, upper arms, wrists, fingers), was used to better interpret the functional relevance of the cortical gradient maps. To identify brain regions recruited for the different movements, we performed one-sample t-tests (*versus* rest) at the group level (Figs. 1d and S5). All movements elicited bilateral activations, primarily located in sensory and motor regions. As expected, activations followed a latero-medial organization, consistent with findings from previous studies (Gordon et al., 2023; Ma et al., 2022; Zeharia et al., 2015). Specifically, jaw movements primarily recruited the lateral part of the sensorimotor cortex, whereas forearm and upper arm movements engaged regions closer to the midline. Wrists and fingers, instead, activated an area situated between these lateral and medial regions. More specifically, the winner-take-all analysis emphasized that the cluster corresponding to finger movements was encircled by the cluster associated with wrist movements.

### 3.3 Delineating spinal segments using cerebro-spinal functional connectivity

To further investigate cerebro-spinal sensorimotor connections and validate their organized nature through a data-driven approach, we then conducted a FC-based parcellation analysis (Figs. 3a, S6a). Our goal was to determine whether distinct functional levels of the spinal cord could be delineated based on their interaction with the brain, specifically its cortical sensorimotor regions. For each participant, matrices representing the degree of similarity in functional connectivity profiles with the sensorimotor iCAP network were computed for all spinal cord voxels. These similarity matrices demonstrated substantial consistency across participants (Fig. S6b, 0.46 ± 0.06, mean correlation across voxels and participants ± standard deviation).

**Figure 3.**
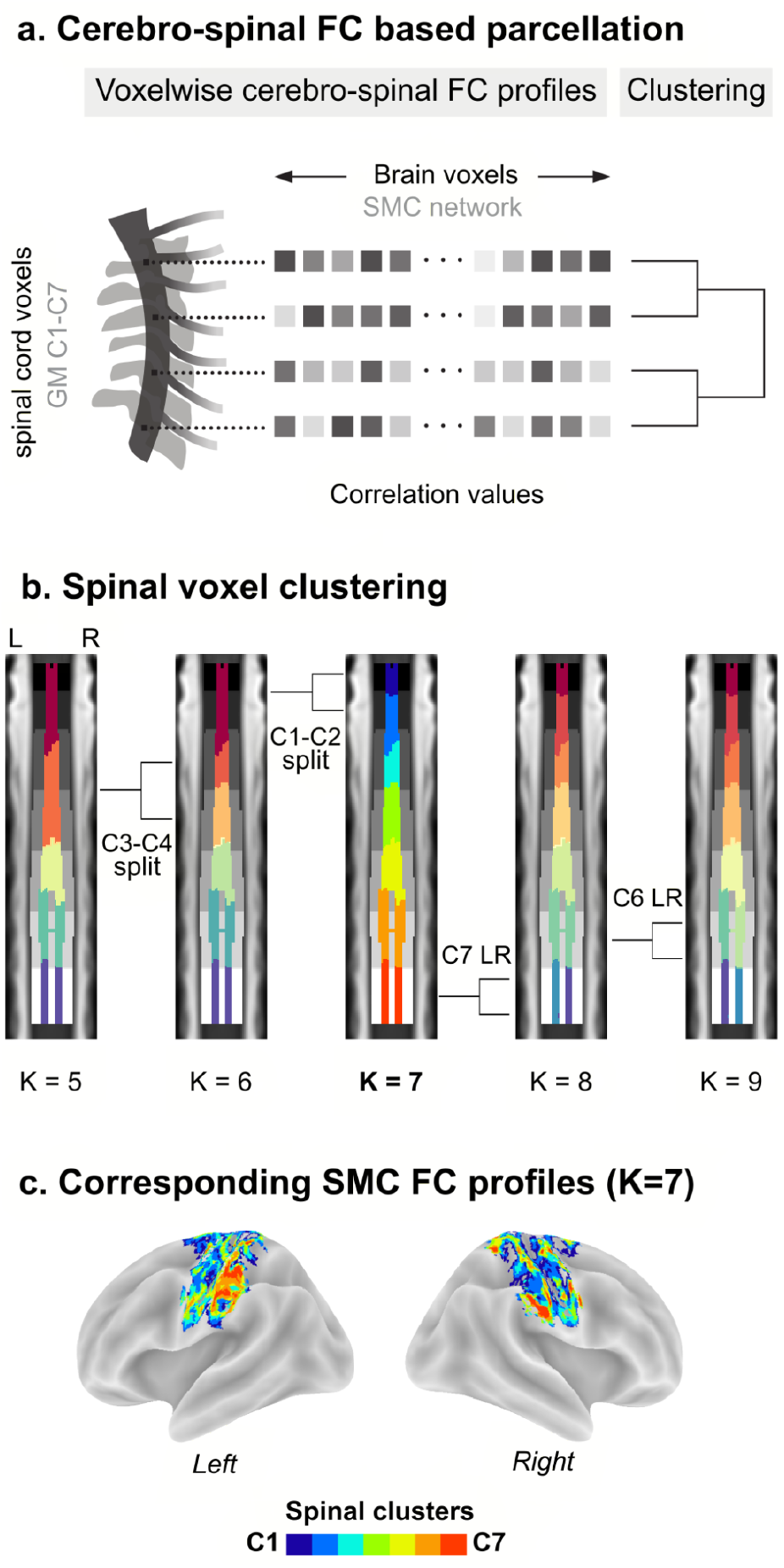
Clustering of spinal cord voxels based on their functional connectivity with the SMC network. **a**. Schematic representation of the FC-based parcellation analysis. For each voxel within the spinal cord (gray matter from C1 to C7), we computed its brain FC profile, as the correlation with each voxel within the cortical SMC network (correlation values represented by boxes with different gray intensities). Then, a hierarchical clustering is used to cluster spinal voxels based on the similarity of their FC profiles with the SMC (Fig. S6a). **b**. Clustering of spinal cord voxels for number of clusters (K) ranging from 5 to 9. Cluster indices are mapped on the spinal cord, with one color per cluster. The central panel shows spinal cord clusters for K = 7. Given the high Dice coefficient between this functionally derived parcellation and spinal segmental levels (Dice = 0.84 ± 0.08, mean across levels ± standard deviation), each label is matched to a spinal level, using the same color scheme as Figure 2. For completeness, we also present cluster maps for neighboring K values, using another colormap to enhance clarity. For each K, a coronal view of the PAM50-T2w template is used as background, cluster indices are depicted with different colors, and spinal levels (Frostell et al., 2016) are shown in grayscale. **c**. Corresponding cerebral signatures for the seven clusters, presented as winner-take-all maps for both hemispheres.

Using this mean similarity matrix as input, we deployed a hierarchical clustering algorithm to explore connectional similarities across spinal cord voxels (Fig. S6a). Adjusting the dendrogram’s cutting point revealed distinct yet related parcellations (Fig. 3b). Our objective of delineating spinal functional levels was achieved by cutting the resulting dendrogram to identify seven (number of clusters denoted as K) spinal cord clusters (central panel of Fig. 3b, Fig. S6a). These functionally delineated clusters varied in size (number of voxels: C1 = 675, C2 = 1013, C3 = 868, C4 = 1436, C5 = 1382, C6 = 1993, C7 = 1633) and were arranged along the spinal cord in a rostro-caudal manner. Importantly, both their spatial extent and localization closely mirrored those of the spinal segmental levels (Dice = 0.84 ± 0.08, mean across levels ± standard deviation) (Frostell et al., 2016). As anticipated, the cerebral functional fingerprints of these seven clusters (Fig. 3c) largely paralleled the gradients derived from the previous analysis (Fig. 2). For other numbers of clusters (Fig. 3b), cutting the dendrogram below K = 7 (*i*.*e*., values lower than the number of segmental levels) resulted in merged clusters (at K = 5, one cluster corresponded to both C1 and C2 spinal segmental levels and another matched both C3 and C4 levels, the latter splitting at K = 6). Above K = 7, within-cluster divisions started to emerge, with the clusters localized at the C7 spinal segmental level dividing into left and right components for K = 8, followed by two clusters at the C6 level for K = 9. The reproducibility of the seven-cluster parcellation was assessed by determining how frequently each of the clusters identified from the group mean were replicated across the 31 participants. We found that all seven clusters could be identified in individual clusterings (Fig. S6c, number of individual parcellations matching each group cluster: C1 = 27, C2 = 18, C3 = 22, C4 = 24, C5 = 25, C6 = 27, C7 = 25). However, individual clusters exhibited a broader extent than the group-level ones, and, for levels C5 to C7, they were more pronounced on the right side.

## 4. Discussion

The investigation of brain and spinal cord somatotopy has traditionally been approached independently, with task-based analyses prevailing in research endeavors. In this study, we proposed a novel approach based on simultaneous neuroimaging of the brain and cervical spinal cord and harnessing the coordinated variations of their sensorimotor spontaneous activities to map large-scale CNS somatotopy. Through two complementary cerebro-spinal functional connectivity analyses, we found compelling evidence of somatotopically organized interactions between these two structures. Our findings emphasize the richness of cerebro-spinal activity, even in the absence of a task, and represent, to the best of our knowledge, the first neuroimaging evidence of large-scale cerebro-spinal somatotopic organization.

First, we employed a data-driven approach, the iCAP framework (Karahanoğlu & Van De Ville, 2015), to isolate sensorimotor networks specific to our population in the brain and spinal cord. Both brain and spinal cord sensorimotor networks extended bilaterally and across somatosensory and motor regions, underscoring the inherent bilateral somatomotor coupling across various levels of the sensorimotor system (Matyas et al., 2010). In the spinal cord, we delineated robust rostro-caudal functional networks corresponding to C1 to C7 segmental levels (Frostell et al., 2016). Drawing upon the fact that nerves from distinct segments project to different body parts (Keegan & Garrett, 1948; Schirmer et al., 2011), we then leveraged segment-wise signals to elucidate brain somatotopy. Analysis of their FC with the SMC revealed a cortical somatotopic gradient, with distinct segments in the spinal cord being preferentially connected with certain regions in the cortex. Crucially, these spinal cord-derived maps of cortical somatotopy were consistent with prior functional and anatomical knowledge. In particular, comparing these maps to task-related maps of body movements (*i*.*e*., jaw, upper arms, forearms, wrists and fingers movements Ma et al., 2022) underscored that spinal cord segments maintain functional connections to their functional homologue areas in the SMC, even during rest.

C3, for instance, was predominantly represented in the lateral part of the precentral and postcentral gyrus, mirroring the activity associated with jaw movements. It is noteworthy that the muscles of the jaw are innervated by cranial – rather than cervical – nerves. Nonetheless, the specific jaw movements examined here (*i*.*e*., biting and twisting), may elicit co-activations of neck muscles, including the infrahyoid and longus capitis muscles, which are innervated by the C3 myotome (Kendall et al., 2005). In addition, C3, through the great auricular nerve, is also involved in innervating the lower portion of the jaw (Ladak et al., 2014). Anastomoses (*i*.*e*., interconnections between nerves), reported between C3 and cranial nerves innervating the face (Shoja et al., 2014), may also contribute to this functional relationship.

In contrast to the lateral representation observed for the C3 level, segments from C4 to C7, which correspond to body territories extending from the shoulders to the fingers, exhibited a more medial distribution within the cortex. Patterns derived from the signals of C4 notably had a cluster appearing closer to the midline, with those related to C6/C7 forming a large cluster extending below. As for task-related activations, upper arms and forearms movements predominantly activated medial regions of the SMC, while movements of the wrists and fingers evoked activity positioned roughly between those associated with jaw and arms movements. This arrangement is consistent with the localization of myotomes and dermatomes corresponding to C4 (shoulders and upper part of the arms) and C6/C7 (forearm to fingers, respectively) (Keegan & Garrett, 1948; Schirmer et al., 2011). A noteworthy observation was the central position of the C7 map within the precentral and postcentral gyrus, surrounded by the C6 and C4/C5 maps. This concentric arrangement, extending from a large representation of the fingers (*i*.*e*., C7 map) at the center to the proximal part of the upper limbs at the periphery, diverged from the continuous, linear somatotopy depicted in textbook homunculi (Penfield & Boldrey, 1937). Analogous concentric representations have been previously reported in the precentral gyrus of humans (Gordon et al., 2023; Meier et al., 2008) and monkeys (Kwan et al., 1978; Park et al., 2001; Wong et al., 1978). To some extent, this pattern was also visible in the winner-take-all maps derived from the task-based activation maps included in this study, as the cluster of the fingers observed in the precentral and postcentral was encircled by a region associated with wrist movements. Hence, our findings complement existing research by suggesting a concentric somatotopic organization across sensory and motor cortices, which does not appear to be exclusive to task-related brain activity, but is also consistently upheld during rest.

While moving or stimulating the jaw and arms is something that can, to a certain extent, be achieved in an MRI scanner, manipulating the neck and scalp is more challenging (accessibility, motion, *etc*.). As C1 and C2 spinal segments primarily innervate myotomes of the neck (Kemp et al., 2011) and dermatomes of the scalp and ear (Ladak et al., 2014), we could not match the corresponding maps to task-related activations. It was however interesting to see that these two levels were largely mapped to the PMc. These results align with anatomical tracing studies in monkeys indicating that corticospinal projections from the PMc primarily terminate in upper cervical segments, with few terminating caudally (He et al., 1993; Morecraft et al., 2019).

The large-scale somatotopic organization of cerebro-spinal functional connections was further corroborated by an additional data-driven analysis, demonstrating the retrieval of spinal cord segments based on the FC profiles of their voxels with the SMC. Employing a hierarchical clustering scheme (Johnson, 1967) offered us the advantage of flexible parcellation, enabling exploration across multiple levels of linkage and ensuring inherent hierarchical consistency in clustering solutions (Eickhoff et al., 2015). Thus, we could explore both the target level of granularity (*i*.*e*., 7, matching the number of spinal segments in our dataset (Frostell et al., 2016) and neighboring solutions. While extracting seven components efficiently captured spinal functional levels based on their shared organization with the brain, reducing the number of clusters led to merged segments. Instead, increasing granularity resulted in the division of C7 and C6 spinal levels into left and right components. This latter observation may reflect variation in the lateralization of the upper body parts. Specifically, the hands, predominantly innervated by caudal cervical segments, necessitate strong lateralization for unilateral goal-directed actions. Conversely, rostral segments innervate regions that do not require the same degree of fine motor control, such as those governing the neck and face. Interestingly, this rostro-caudal difference in lateralization echoed the patterns obtained when assessing the inter-participant reproducibility of the parcellation. Notably, participants’ components corresponding to lower cervical segments were more prominently localized on the right side, a phenomenon potentially associated with the right-handed dominance in our study population.

These combined analyses shed new light on the intricate relationships between the brain and cervical spinal cord, even during rest, adding depth to earlier work that showed organized patterns of cerebro-spinal FC (Vahdat et al., 2020), albeit without specific investigation into somatotopy. Previous studies have independently reported organization in both the brain and spinal cord, emphasizing local processing of somatosensory inputs and motor outputs between homologous regions (*i*.*e*., regions associated with the same body parts across the two hemispheres or the two spinal hemicords). For instance, brain studies have demonstrated that hand representation in the SMC exhibits stronger FC with the contralateral hand region than with other body parts (*e*.*g*., face and foot Gordon et al., 2016; Thomas et al., 2021). Evidence also suggests that within-hemisphere brain somatotopy can be observed during resting-state (Long et al., 2014). Similarly, in the spinal cord, FC has been shown to be higher within the same segmental level compared to FC between segmental levels (Kinany et al., 2020; Kong et al., 2014; Landelle et al., 2023). These brain and spinal cord studies provided compelling evidence that the strength of FC at rest aligns with the functional relevance of adjacent body territories, exhibiting stronger FC for areas requiring higher coordination and interaction. Our findings further contribute by suggesting that the sensorimotor system at rest is not merely divided into local networks; instead, these analogous organizational patterns seem to represent a general feature of the sensorimotor hierarchy. Likely mediated through descending (motor) and ascending (sensory) pathways (Usuda et al., 2022), this large-scale organization allows for the unified processing of information from specific body parts.

The results of the present study underscore the richness of spontaneous activity in the CNS and its potential to unveil the large-scale functional architecture of the somatosensory hierarchy with minimal experimental burden. Such investigations can offer a valuable lens into the processes underlying CNS-body interaction, with broad implications spanning fundamental neuroscience and clinical practice. Plastic changes in the sensorimotor system are fundamental to continuously shape the neural circuits that allow us to acquire new skills from infancy to adulthood, to compensate for aging, and to adjust to injuries and conditions affecting the CNS or the body. For instance, the loss of somatosensory inputs from a body region following amputation (Muret & Makin, 2021) or spinal cord injury (Freund et al., 2013) can lead to remodeling of the sensorimotor functional architecture from the periphery to the cortex. Likewise, alterations in body movement and perception resulting from normal aging and age-related neurodegenerative diseases like Parkinson’s disease, can impact spontaneous neural activity related to the affected body parts at both brain (Caspers et al., 2021; Landelle et al., 2020) and spinal cord levels (Landelle et al., 2023). These alterations likely result from the complex interplay of bottom-up and top-down processes, indicating a systemic reorganization rather than isolated local changes. Consequently, leveraging cerebro-spinal fMRI to probe somatotopy holds significant promise for elucidating these neuroplastic mechanisms on a comprehensive scale, thus advancing our understanding of CNS-body interactions.

## Data availability

Data and other materials that directly support the main conclusions of the present study are available in the main manuscript or in Supplementary information.

## Acknowledgments

This work was performed at the McConnell Brain Imaging Centre, The Neuro (McGill University, Montreal Canada). We thank David Costa, Ronaldo Lopez and Soheil Mollamohseni Quichani for their help in data acquisition and all participants who were recruited for this study.

C.L was supported by Fonds de Recherche du Québec—Santé (FRQ-S) and the Canadian Institutes of Health Research (CIHR). N.K. and D.V. were supported by funds from the Swiss National Science Foundation (Project No. 205321_207493). J.D. has received grants to support that study from Fondation Courtois, the Natural Sciences and Engineering Research Council of Canada (NSERC, Grant No. 248074); Brain Canada Platform (Grant No. 255934) and Healthy Brain for Healthy Lives (HBHL, Grant No. 2473633). The funders had no role in study design, data collection and analysis, decision to publish, or preparation of the manuscript.

## Author contributions

Conceptualization - B.DL., C.L., J.D., and N.K

Data acquisition - C.L., and ND.M

Methodology - B.DL., C.L., N.K., and D.VDV.,

Formal Analysis - C.L., N.K., and ND.M.

Writing – Original Draft, C.L., and N.K.

Writing - C.L., and N.K.

Review and Editing - all the authors.

Funding Acquisition - J.D.

Supervision - J.D., and D.VDV.

