## Supplementary information for "Cerebro-spinal somatotopic organization uncovered through functional connectivity mapping"

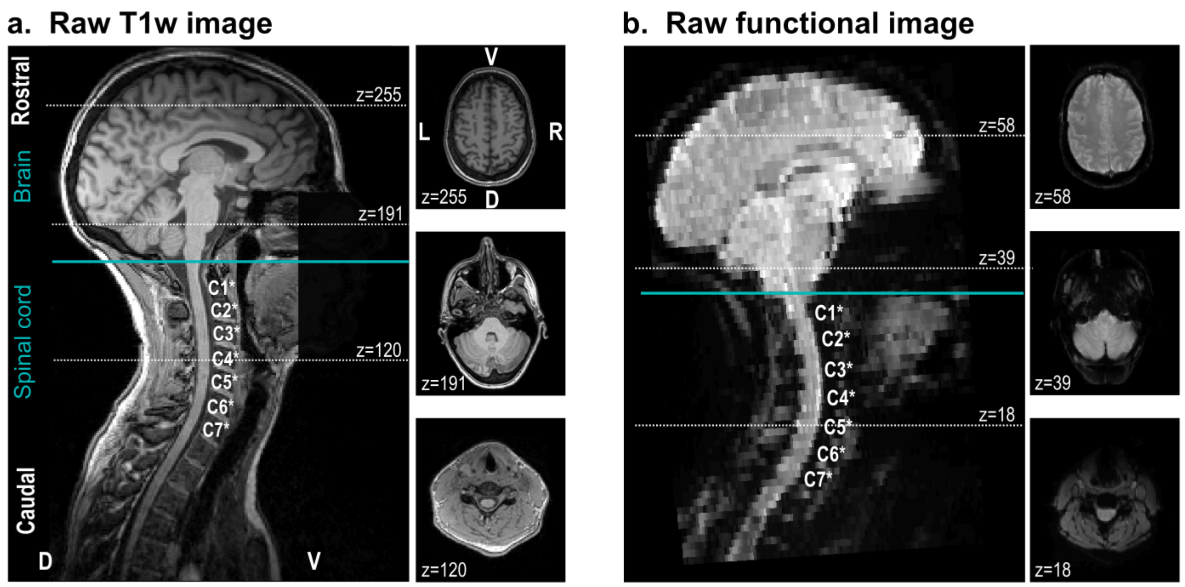

**Figure S1 | Data quality.** Raw anatomical (a.) and functional (b.) images for an example participant. The field of view included the whole brain and cervical spinal cord. D: dorsal; V: ventral; L: left; R: right. Images are in the participant space. Labels from C1* to C7* refer to vertebral levels.

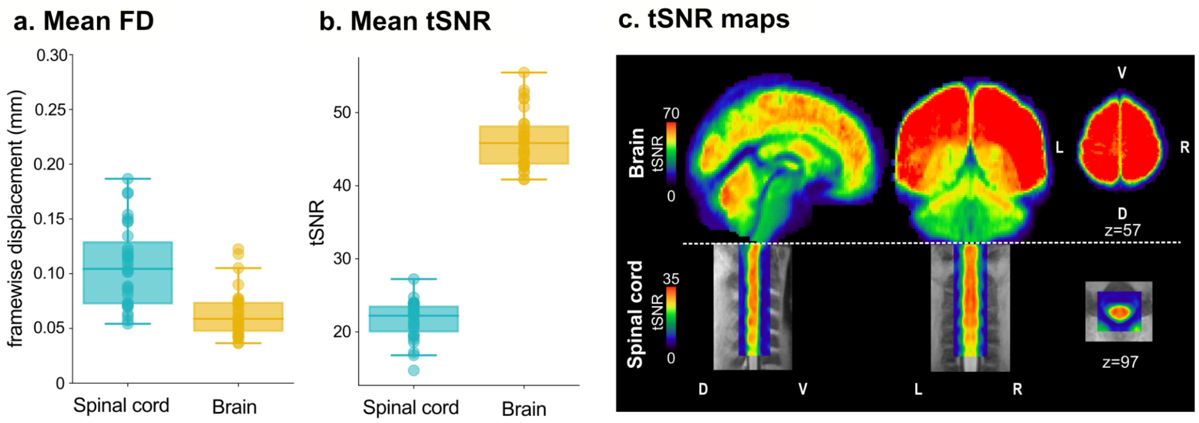

**Figure S2 | Quality check metrics. a-b.** Boxplots of the tSNR and the framewise displacement (FD) for both spinal cord (blue) and brain (yellow) masks. Each box extends from the 25th to the 75th percentile of the group's distribution and the medians are represented by the horizontal line inside the box. The vertical extending lines denote the extreme values within 1.5 interquartile range. **c.** Average temporal signal-to-noise ratio (tSNR) maps for both the brain and the spinal cord are presented. Different scales are used to better visualize variations within each region. D: dorsal; V: ventral; L: left; R: right.

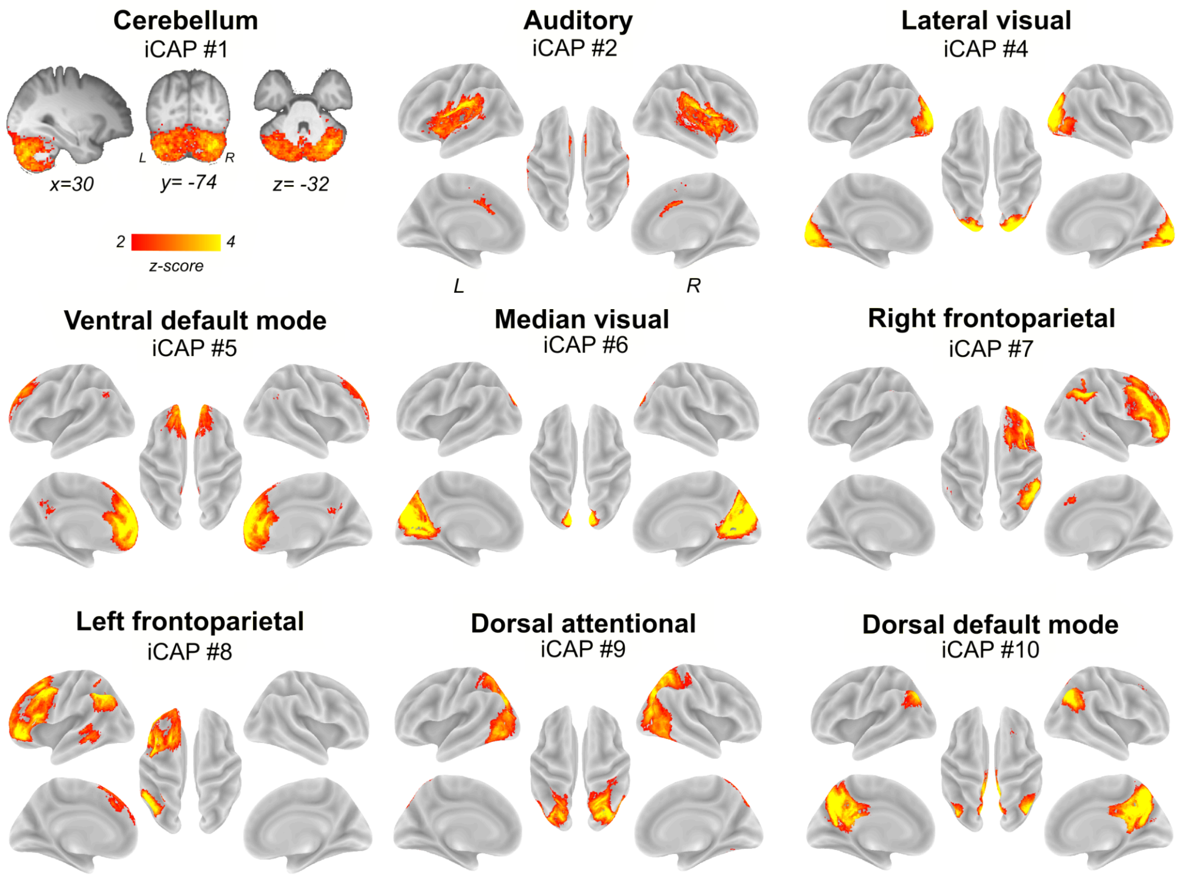

**Figure S3 | iCAPs resting-state brain networks**. Spatial patterns for the 10 innovation-driven coactivation patterns (iCAPs). Note that the map for iCAP #3 (sensorimotor cortical network) is available in the main text, (Fig 1A). iCAP #1 is overlaid on the average participants' T1w anatomical image (x,y,z coordinates in MNI space). L: left; R: right.

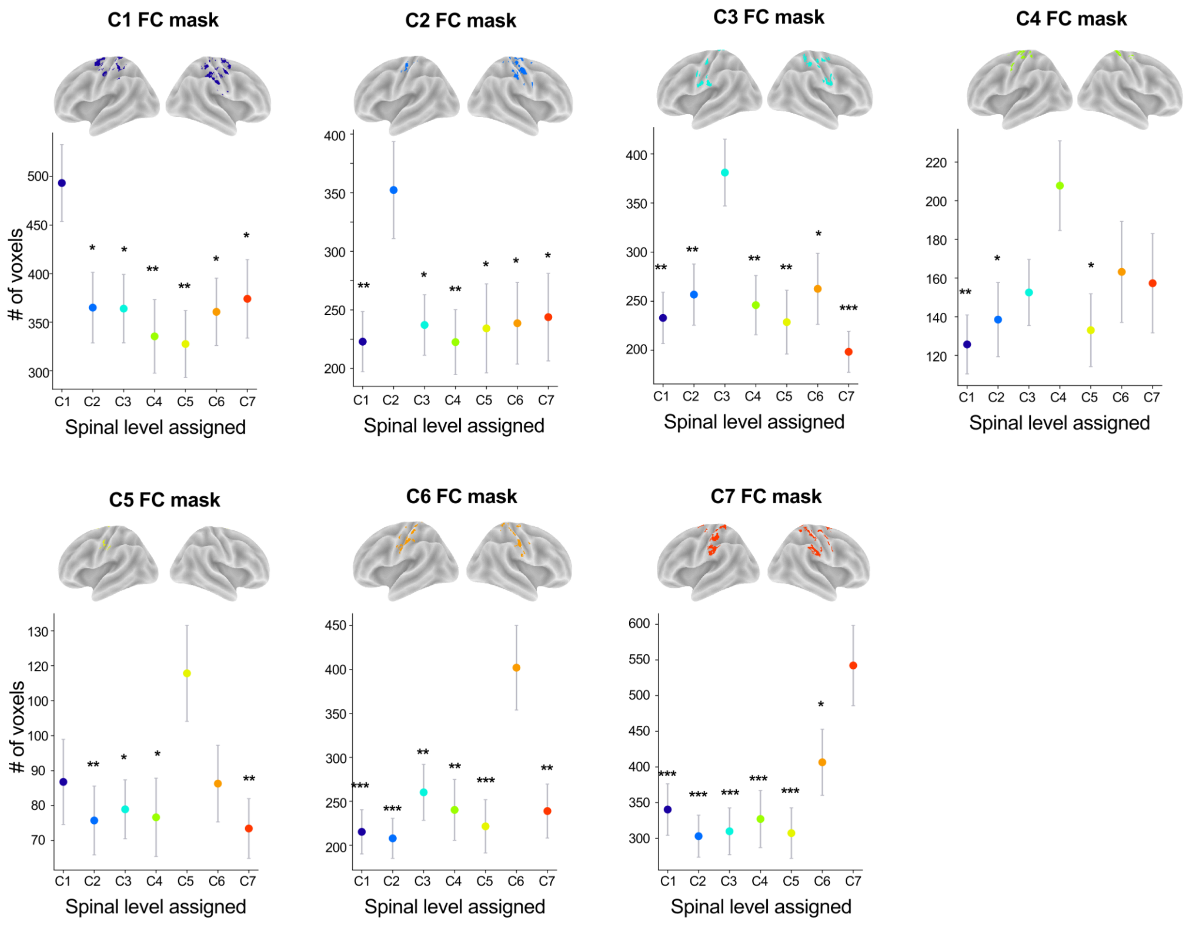

**Figure S4 | Reproducibility of the winner-take-all maps for the seed-to-voxels analyses.** The winner-take-all maps were obtained for each individual and the mean number of voxels assigned to each spinal cord level was calculated within each group-level FC map. For each FC mask and level assigned, dots indicate the mean number of voxels and error bars to the standard error of the mean. Reported statistics correspond to the fixed effect of linear mixed models (see details in Table S1).* p<0.05, ** p<0.01; ***p<0.001

| A. C1 FC mask | B. B. C2 FC mask | C. C. C3 mask | D. C4 FC mask |
|---|---|---|---|
| C1 vs. C2: t=2.30; p=0.022 | C2 vs. C1: t=2.61; p=0.0095 | C3 vs. C1: t=3.15; p=0.0018 | C4 vs. C1:t=2.61; p=0.009 |
| C1 vs. C3: t=2.32; p=0.021 | C2 vs. C3: t=2.32; p=0.021 | C3 vs. C2: t=2.64; p=0.0084 | C4 vs. C2: t=2.21; p=0.028 |
| C1 vs. C4: t=2.83; p=0.0050 | C2 vs. C4: t=2.62; p=0.0090 | C3 vs. C4: t=2.87; p=0.0045 | C4 vs. C3: t=1.76; p=0.079 |
| C1 vs. C5: t=2.97; p=0.0033 | C2 vs. C5: t=2.39; p=0.018 | C3 vs. C5: t=3.24; p=0.0013 | C4 vs. C5: t=2.39; p=0.018 |
| C1 vs. C6: t=2.38; p=0.018 | C2 vs. C6: t=2.30; p=0.022 | C3 vs. C6: t=2.52; p=0.012 | C4 vs. C6: t=1.42; p=0.16 |
| C1 vs. C7: t=2.14; p=0.033 | C2 vs. C7: t=2.19; p=0.029 | C3 vs. C7: t=3.88; p<0.001 | C4 vs. C7: t=1.61; p=0.11 |

| E. C5 FC mask | F. C6 mask | G. C7 mask |
|---|---|---|
| C5 vs. C1: t=1.94; p=0.054 | C6 vs. C1: t=3.76; p<0.001 | C7 vs. C1: t=3.37; p<0.001 |
| C5 vs. C2: t=2.62; p=0.0092 | C6 vs. C2: t=3.91; p<0.001 | C7 vs. C2: t=3.99; p<0.001 |
| C5 vs. C3: t=2.43; p=0.016 | C6 vs. C3: t=2.86; p=0.0046 | C7 vs. C3: t=3.88; p<0.001 |
| C5 vs. C4: t=2.57; p=0.011 | C6 vs. C4: t=3.26; p=0.0012 | C7 vs. C4: t=3.59; p<0.001 |
| C5 vs. C6: t=2.97; p=0.050 | C6 vs. C5: t=3.64; p<0.001 | C7 vs. C5: t=3.92; p<0.001 |
| C5 vs. C7: t=2.77; p=0.0060 | C6 vs. C7: t=3.29; p=0.0012 | C7 vs. C6: t=3.26; p=0.024 |

**Table S1 | Reproducibility of the winner-take-all maps for the seed-to-voxels analyses (statistical results).** The winner-take-all maps were obtained for each individual and the mean number of voxels assigned to each spinal cord level was calculated within each group-level FC map. The fixed effect of the seven linear mixed models is reported, the degree of freedom is 210 for all tests, the t and p values are reported for each fixed-effects.

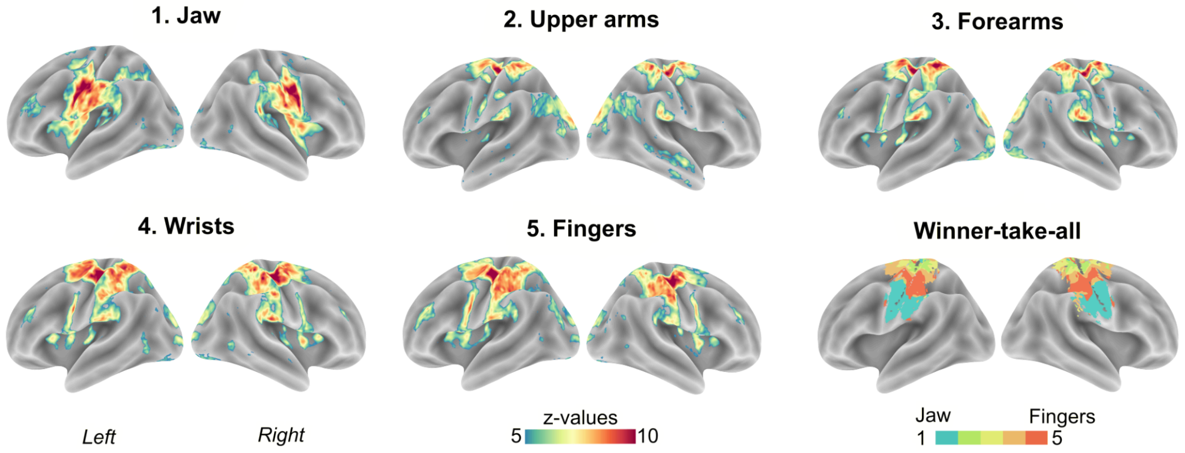

**Figure S5 | Movement related brain activity patterns.** Group-level (61 participants) statistical Z-maps were obtained with one-sample t-test contrasting each movement conditions with the rest period. Statistical maps were thresholded at Z = 5 (p<0.000001 uncorrected). The bottom right panel shows winner-take-all maps for these five conditions, in our sensorimotor iCAP mask. Task-related data were derived from a publicly available dataset (Ma et al., 2022).

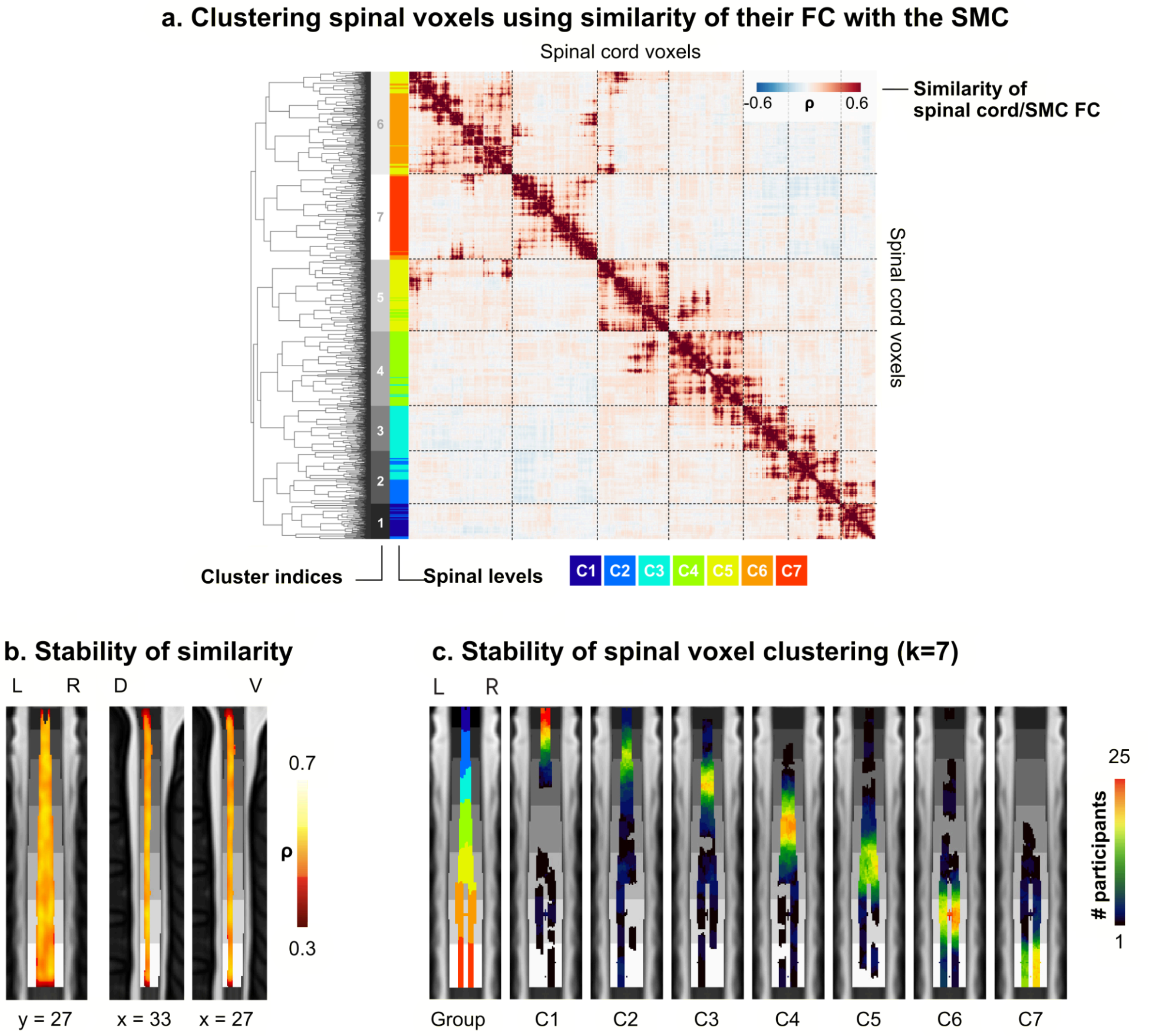

**Figure S6 | FC-based parcellation analyses. a.** For each pair of spinal cord voxels, the matrix illustrates the average similarity in functional connectivity (FC) profiles with the sensorimotor cortex (SMC) iCAP network. Colors denote the strength of correlations between these profiles (red for positive correlation, blue for negative). This matrix underwent hierarchical clustering to explore brain connectional similarities across spinal cord voxels, resulting in the dendrogram shown on the left. Rows and columns were reorganized based on these patterns of connectional similarities. With the aim of delineating spinal functional levels, we cut the resulting dendrogram to identify seven spinal cord clusters (*i.e.*, K = 7), with corresponding cluster indices indicated by gray blocks. To assess the agreement between these functionally-derived parcellation and the functional organization of the spinal cord, each voxel is also colored based on its spinal segmental level. **b.** Inter-individual stability of similarity matrices. Maps illustrate the average correlation between the similarity profiles (*i.e.*, similarity in FC profiles with the sensorimotor iCAP network for each pair of spinal cord voxels), across all participants. The left panel shows a coronal view, while the two right panels depict left (x = 33) and right (x = 27) sagittal views. **c.** Stability of the seven-cluster parcellation. We determined how frequently each of the seven clusters identified from the group mean (left panel) were replicated across participants. The right panels contain the distribution maps for each cluster, which was assigned to the corresponding spinal level. The PAM50-T2w template is used as background, and spinal levels (Frostell et al., 2016) are shown in grayscale. D: dorsal; V: ventral; L: left; R: right.
